# Location of the TEMPO moiety of TEMPO-PC in phosphatidylcholine bilayers is membrane phase-dependent

**DOI:** 10.1101/2022.03.18.484922

**Authors:** Seonghoon Kim, Changbong Hyeon

## Abstract

The (2,2,6,6-tetramethylpiperidin-1-yl)oxyl (TEMPO) moiety tethered to the headgroup of phosphatidylcholine (PC) lipid is employed in spin labeling electron paramagnetic resonance (EPR) spectroscopy to probe the water dynamics near lipid bilayer interfaces. Due to its amphiphilic character, however, TEMPO spin label could partition between aqueous and lipid phases, and may even be stabilized in the lipid phase. Accurate assessment of the TEMPO-PC configuration in bilayer membranes is essential for correctly interpreting the data from measurements. Here, we carry out all-atom molecular dynamics (MD) simulations of TEMPO-PC probe in single-component lipid bilayers at varying temperatures, using two standard MD force fields. We find that for DPPC membrane whose gel-to-fluid lipid phase transition occurs at 314 K, TEMPO is stabilized above (below) the bilayer interface if the membrane is in the gel (fluid) phase. For bilayers made of unsaturated lipids, DOPC and POPC, which adopt the fluid phase at ambient temperature, TEMPO is unequivocally stabilized inside the bilayers. Our finding of membrane phase-dependent positioning of TEMPO moiety highlights the importance of assessing the packing order and fluidity of lipids under a given measurement condition.

## Introduction

TEMPO is a six-membered cyclic nitroxide radical shielded by four methyl groups. Tethered at various positions along the hydrocarbon tail in the form of *n*-doxyl PC lipids, fluorophores or nitroxide spin labels including TEMPO have been employed to measure the penetration depth of solvent molecules and the flexibility of lipid tail by analyzing homogeneous line broadening of EPR spectra^1–4^ and fluorescence quenching. ^5^ Meanwhile, TEMPO-PC, a molecular construct in which TEMPO molecule is tethered to the choline group of PC, has been used to probe the water/lipid interfacial molecular environment^6–8^ with an estimate based on molecular model that TEMPO is positioned at 5 Å above the bilayer interface. ^9–11^

The location of TEMPO moiety in PC bilayers is, however, not entirely clear and remains controversial in the recent literature. A series of MD simulation and fluorescence quenching-based studies have been showing that PC headgroup-labeled probes, e.g., 1,6-diphenyl-1,3,5-hexatriene (DHP), 7-nitrobenz-2-oxa-1,3-diazol-4-yl (NBD), and TEMPO reside in the acyl chain region. ^12–17^ In contrast, it has been argued^18^ that the headgroup-labeled spin probes residing in the aqueous phase, protruded at ∼ 5 Å above the bilayer interface, are evidenced by EPR spectra.

Here, we investigate the location of TEMPO spin label of TEMPO-PC in PC bilayers by carrying out a set of careful simulations. We first perform brute-force MD simulations at different temperatures to get rough ideas on the location of TEMPO moiety in DPPC bilayers, and next determine thermodynamically more reliable location of TEMPO by calculating the potential of mean forces (PMFs) via the umbrella sampling method. PMFs of TEMPO moiety in *unsaturated* PC bilayers (DOPC, POPC) are also calculated. The PMFs obtained with three lipid types at varying temperatures using two different MD force fields allow us to draw a general conclusion that the location of TEMPO moiety is membrane phase-dependent. To be specific, if the bilayers are in fluid phase, TEMPO-PC adopts a configuration that TEMPO group is stabilized inside bilayers (TEMPO-in configuration), whereas if the bilayers are in gel phase, TEMPO-PC adopts the TEMPO-out configuration in which the TEMPO group is stabilized above the lipid-water interface.

## RESULTS

### TEMPO-PC in DPPC bilayers at varying temperatures

We first carry out a series of 500 ns brute-force MD simulations of a fully hydrated DPPC bilayer system containing ∼ 3 mol% TEMPO-PC at varying temperatures by using the CHARMM36 force field. The snapshots of simulation visualize how the configurations of lipid bilayer and TEMPO-PC change with temperature (Fig. 1A). At high temperatures (*T* = 320, 330 K), the bilayer membrane adopt fluid phase, and the TEMPO-PC adopts the TEMPO-in configuration, such that TEMPO moiety is stabilized inside the bilayer interface. At low temperatures, especially in the range of 290 K ≤ *T* ≤ 310 K, the acyl chains of lipids adopt ordered configuration, and partial overlaps between the upper and lower leaflets give rise to corrugation in the bilayer surface, reminiscent of the *P*_*β*_ (rippled) phase. TEMPO-in and TEMPO-out configurations coexist in this temperature range, yielding the bimodal distributions in the TEMPO location normal to the membrane. At even lower temperature (*T* = 260 and 270 K), the TEMPO-out configuration becomes dominant (Fig. 1A).

**Figure 1:**
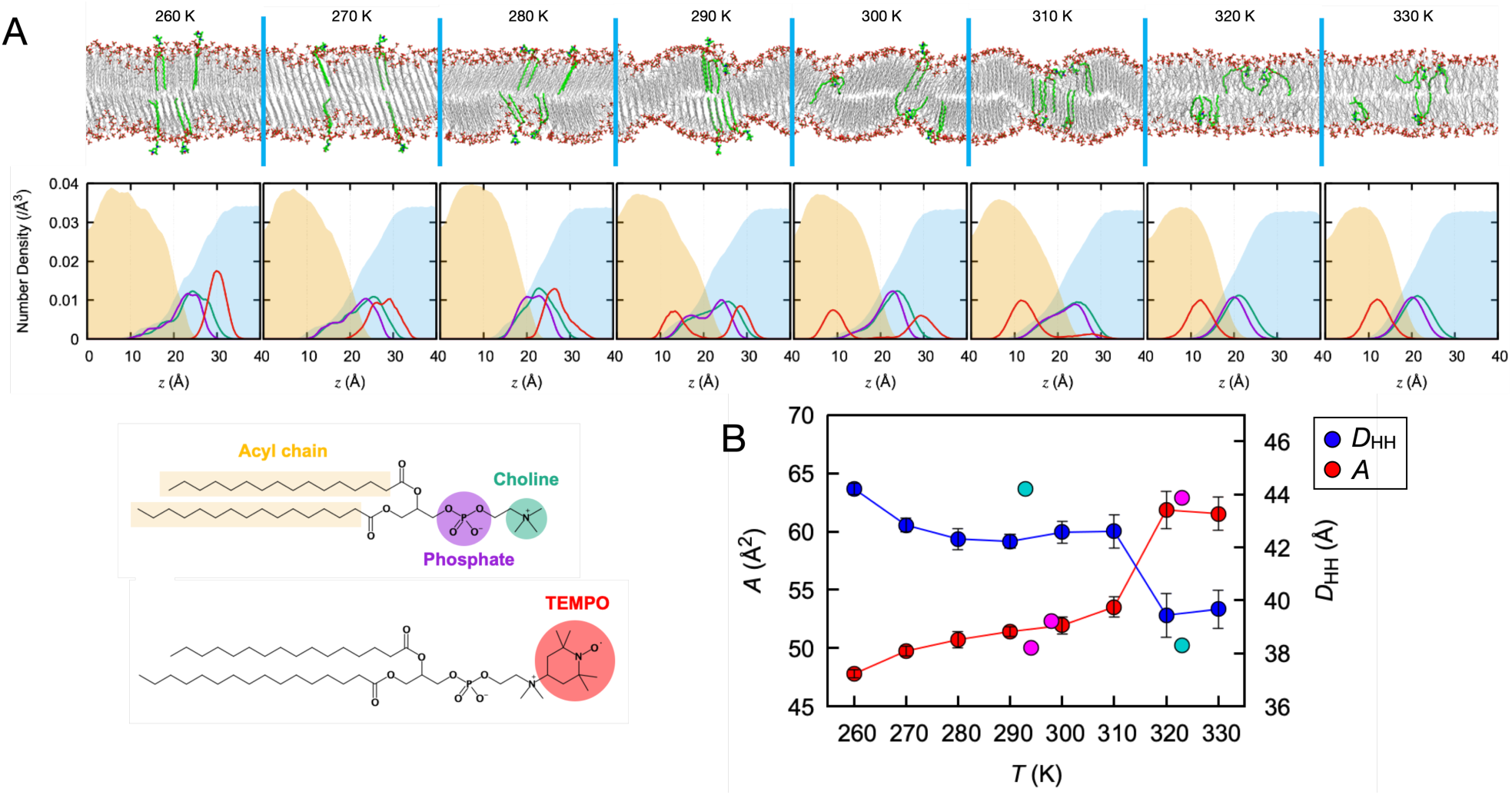
MD simulations of TEMPO-PC in DPPC bilayers at varying temperatures. **A**. Snapshots of TEMPO-PC configurations in DPPC bilayers (first row). The interdigitated state characterized with the overlap between upper and lower leaflets and corrugation in membrane surface are observed at 290 K ≤ *T* ≤ 310 K. The density profiles of the heavy atoms in phosphate (magenta) and choline groups (green) of DPPC and TEMPO group (red) calculated from the simulations are shown in the second row, along with acyl chain in DPPC (filled ivory), and water (filled pale blue). The density of TEMPO moiety (red) is amplified by 5 folds for clarity. **B**. Area per lipid (*A*) and bilayer thickness (*D*_HH_) of DPPC bilayers obtained from MD simulations at varying temperatures (error bars denote the standard deviation) that used the CHARMM36 force field. The filled circles in magenta and cyan represent the measurements^22^ for *A* and *D*_HH_, respectively. The gel-to-fluid phase transition temperature of DPPC membranes is estimated to be *T*_m_ = 314 K from the crossing point of *A* and *D*_HH_.

The two key physical characteristics of the DPPC bilayers, area per lipid (*A*) and bilayer thickness (*D*_HH_), the latter of which measures the average distance between the phosphate groups in the upper and lower leaflets are in reasonable agreement with experimental measurements (filled circles in magenta and cyan in Fig. 1B). From *A*(*T*) and *D*_HH_(*T*), the gel-to-fluid (or *P*_*β*_-to-*L*_*α*_) lipid phase transition temperature (*T*_m_ = 314 K) is correctly identified.^19–24^

### Potential of mean force of TEMPO spin label in DPPC bilayers

In the foregoing result from the brute-force MD simulations, simulation time may not be long enough to give thermodynamically reliable results. The simulations offer only a rough idea of TEMPO-PC configurations. To determine the temperature-dependent configuration of TEMPO-PC in the DPPC bilayers, we carry out the umbrella sampling to calculate the PMF of TEMPO group across the DPPC bilayer using two standard force fields, CHARMM36 and AMBER LIPID17. To avert the possibility that two different TEMPO-PCs in the same leaflet interact with each other and compromise our PMF calculation, we place only a single TEMPO-PC in each leaflet, effectively making a system of DPPC bilayer containing ∼ 1.5 mol% TEMPO-PC.

Because of substantial fluctuations, corrugation of bilayer interface, and the temperature-dependent membrane thickness (Fig.1A), the distance from the bilayer center is not a convenient metric to discuss whether the TEMPO moiety is in the interior or exterior of the bilayer interface. Thus, for straightforward determination of the TEMPO location, we set the average position of phosphate group as the reference penetration depth (Δ*z* = 0) of TEMPO, so that we can easily assess from PMF whether the TEMPO-PC adopts TEMPO-in (Δ*z <* 0) or TEMPO-out (Δ*z >* 0) configuration. In addition, we monitor the minimal distance (*d*_TA_) between the methyl groups of TEMPO and the upper part of hydrocarbon tail (C2-C5 of acyl groups) as another coordinate to characterize the TEMPO-PC configuration. The two dimensional (2D) PMFs calculated as a function of Δ*z* and *d*_TA_ (Fig. 2) demonstrate that the dominant population of TEMPO-PC configuration is formed at Δ*z* = 5 Å and *d*_TA_ ≈ (8−12) Å at *T* = 290 K, and at −10 Å *<* Δ*z <* −5 Å and *d*_TA_ ≈ (3 − 4) Å at *T* = 320 K. The former denotes the TEMPO-out configuration, more specifically, TEMPO moiety is stabilized in the aqueous phase, being excluded from the compactly ordered lipid phase. On the other hand, in the latter configuration (−10 Å *<* Δ*z <* −5 Å and *d*_TA_ ≈ (3 − 4) Å), TEMPO-PC adopts bent configurations in which the TEMPO group is stabilized in the lipid phase.

**Figure 2:**
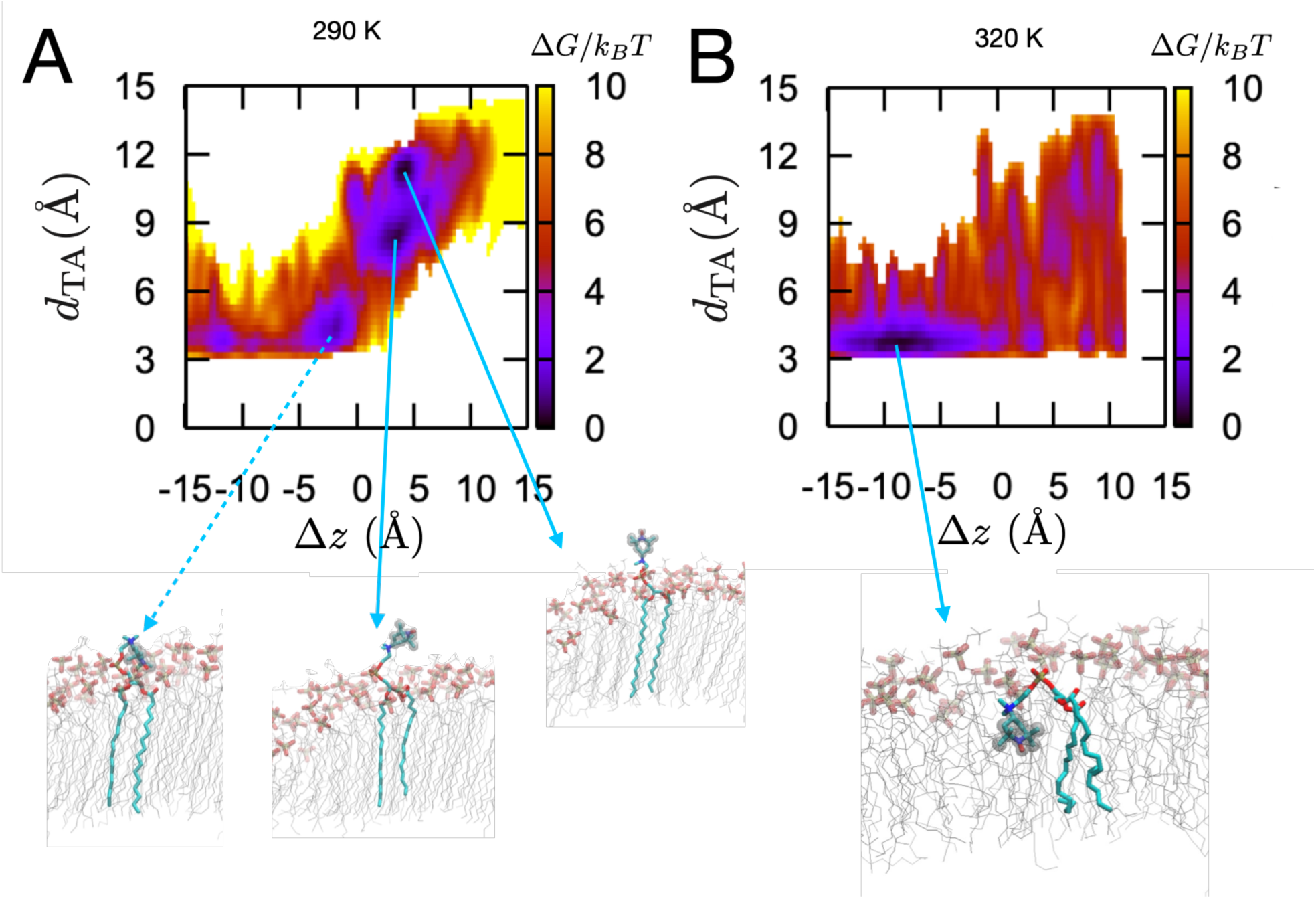
2D PMF of TEMPO spin label in the DPPC bilayers calculated as a function of Δ*z* (height difference between nitroxide oxygen and phosphorus atoms) and *d*_TA_ (minimal distance between any heavy atom of TEMPO and C2-C5 of acyl chain at (A) *T* (= 290 K) *< T*_m_ and *T* (= 320 K) *> T*_m_. Depicted are the snapshots of TEMPO-PC configuration at free energy minima with phosphate groups of DPPC visualizing the lipid-water interface. TEMPO-out and TEMPO-in configurations are dominant at *T < T*_m_ and *T > T*_m_, respectively. The calculation was carried out under AMBER LIPID17.

One dimensional projection of 2D PMF in Fig.2 leads to the 1D PMFs demonstrated in Fig.3. The results obtained using the AMBER LIPID17 force field reconfirm that the TEMPO is stabilized at Δ*z* ∼ 5 Å, i.e., above the bilayer interface of the DPPC membrane at T = 290 K, so that TEMPO-out configuration becomes dominant (Fig. 3A). In contrast, at *T* = 300, 310 and 320 K, TEMPO is stabilized at the penetration depth of −10 Å *<* Δ*z <* −5 Å with free energy bias of ∼ (5 − 7) *k*_*B*_*T* towards the lipid phase (Fig. 3A). The 1D PMFs under CHARMM36 yield a similar tendency except for *T* = 300 K at which the TEMPO-out configuration is favored (Fig. 3B). We note that *T*_m_ estimated for the DPPC bilayers under AMBER LIPID17 is lower than the value under CHARMM36 by ∼ 10 K (compare Fig. S1 with Fig. 1B).

**Figure 3:**
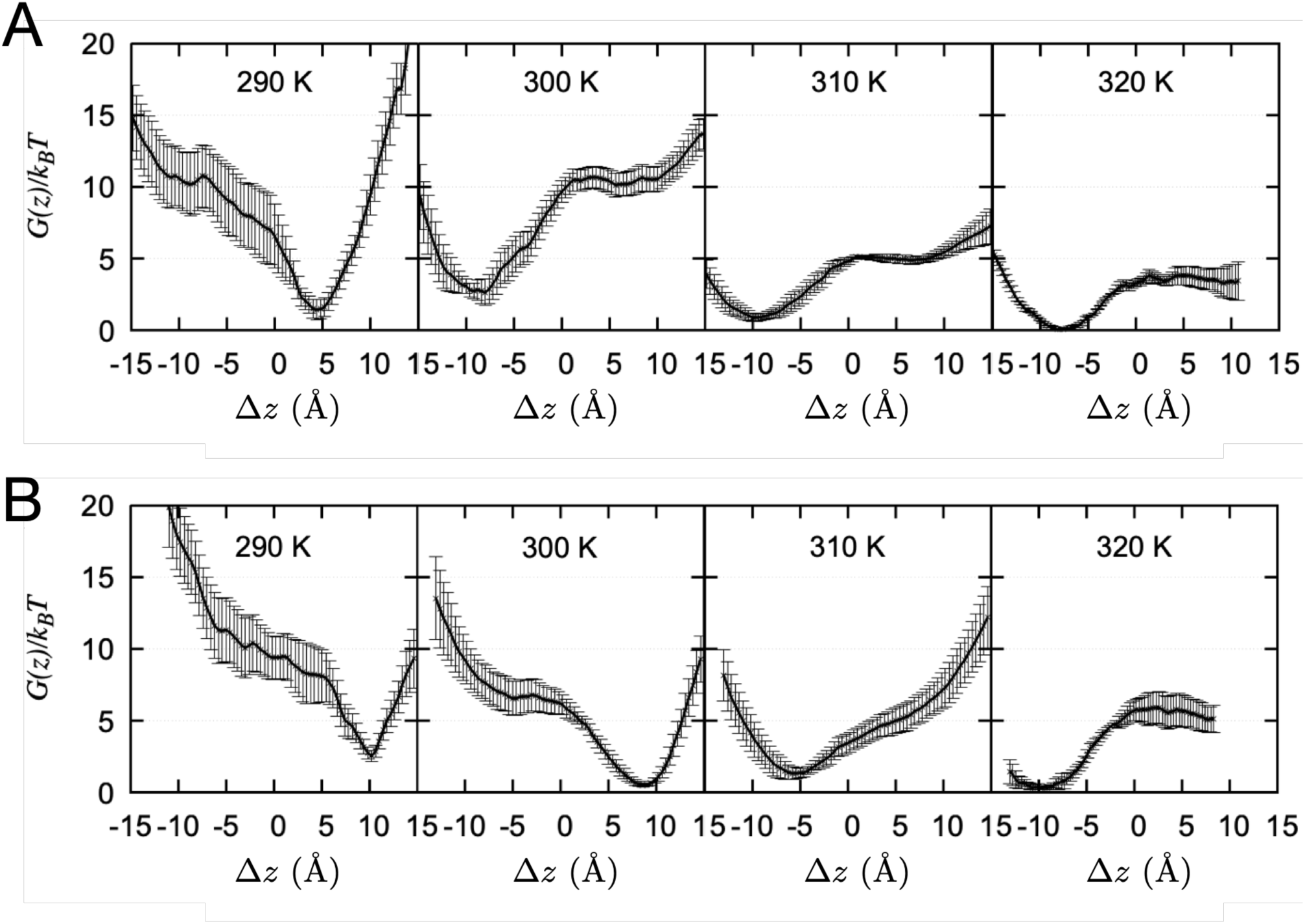
PMFs of TEMPO spin label (nitroxide oxygen) in the DPPC bilayers at various temperatures (error bars are estimated standard error) calculated using the **A**. AMBER Lipid17 and **B**. CHARMM36 force fields.

### PMFs of TEMPO spin-label in unsaturated lipid bilayers

Bilayers formed from unsaturated lipids, such as POPC and DOPC characterized with low lipid phase transition temperature (*T*_m_ −2 ^°^*C* for POPC^25,26^ and *T*_m_ −16.5 ^°^*C* for DOPC^27^), are in the fluid phase at ambient temperature. We repeated the PMF calculation of TEMPO spin label along the Δ*z*-axis in POPC and DOPC membranes at *T* = 310 K using the two force fields (Fig. 4). Besides the variations with lipid types and force fields, the PMFs of TEMPO unequivocally indicate that TEMPO-in configuration is more favorable. The thermodynamically favorable position of TEMPO at −10 ;S Δ*z* ;S −5 Å with the free energy bias of (5−7) *k*_*B*_*T* is consistent with our previous MD simulation study of TEMPO-PC in POPC bilayers modeled with Berger force field.^15^

**Figure 4:**
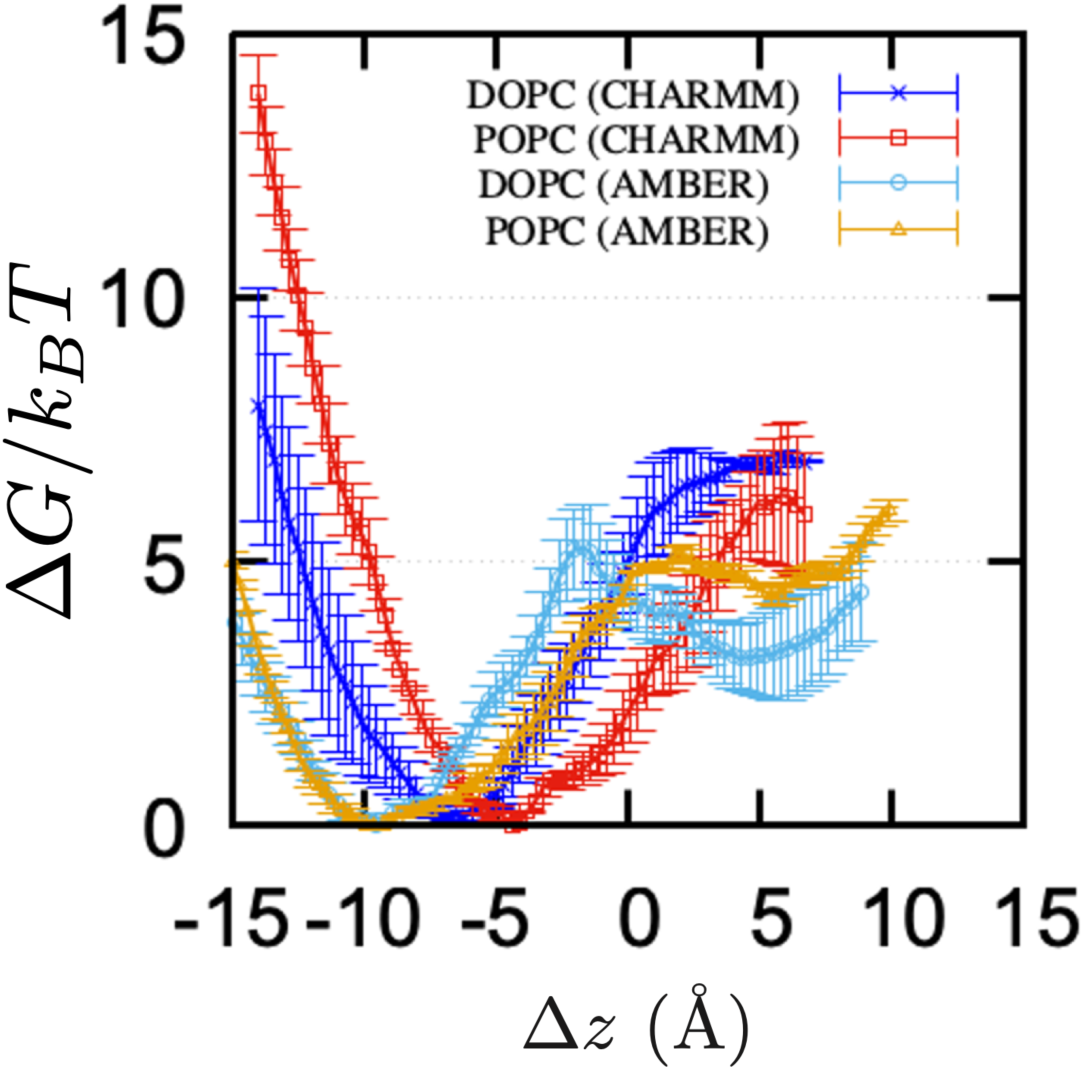
PMFs of TEMPO spin label in unsaturated lipid (POPC and DOPC) bilayers at *T* = 310 K calculated using the CHARMM and AMBER force fields (the error bars denote estimated standard error).

## Discussion

Here, we discuss our key finding of membrane phase-dependent location of TEMPO moiety in light of historical perspective, current observations, and the questions raised over the past years.

1. It has been known from earlier studies in 1970s that unsubstituted TEMPO molecules can partition between fluid hydrophobic regions of lipids and the aqueous region. ^28^ The partition coefficient or the solubility of TEMPO to the lipid phase increases with the fluidity of the lipids, whereas it is excluded from paracrystalline regions (gel phase) of lipid bilayers.^28^ Thus, an empirical relationship between the partition coefficient of TEMPO and the lipid order parameter determined with EPR signals (*A*_*∥*_ outer hyperfine splitting) from TEMPO-labeled fatty acid (say, 5-PC) allows one to determine the fraction of lipid in the fluid-phase membrane, ^28,29^ enabling to construct the phase diagram of phospholipid mixtures. ^30,31^
There may be a certain difference between the molecular behaviors of the unsubstituted TEMPO and the TEMPO moiety attached to PC-headgroup in bilayer membranes. However, the TEMPO-labeled headgroup is flexible and subject to larger thermal fluctuations. It is noteworthy that fluctuations of TEMPO moiety buried inside the membrane at *T > T*_m_ are still larger than the fluctuations of TEMPO residing in the aqueous phase at *T < T*_m_. Fluctuations in TEMPO position increase gradually with temperature (see Fig. S2). In light of the unsubstituted TEMPO partitioning to fluidic lipid phase and its exclusion from gel phase, the membrane phase-dependent behavior of TEMPO-PC configuration, adopting the TEMPO-in (fluid phase) and TEMPO-out (gel phase), is not entirely unexpected.
2. Because TEMPO moiety switches its location from the aqueous phase at low temperature (*T < T*_m_) to the fluidic lipid phase at high temperature (*T > T*_m_), the change in TEMPO mobility at *T* ∼ *T*_m_ would be not as discontinuous as 5-PC and 5-DSA^7,18^ or a hypothetical case of TEMPO being buried inside the bilayers over the entire temperature variation. This in part rationalizes the *absence* of sharp (discontinuous) jump in the EPR-measured rotational diffusion rate and diffusivity of TEMPO across *T*_m_ ^7,18^ (see Fig.S2 as well).
3. The membrane phase-dependent location of TEMPO simultaneously rationalizes both the nitrogen hyperfine coupling constant data (2*A*_*z*_) at *T* = −165 ^*°*^C (Fig. 8b in Ref.^32^) and the oxygen transport parameter at *T* = 25 ^*°*^C (Fig. 9b in Ref.^32^) for *n*-PC and TEMPO-PC in pure POPC bilayers by Subczynski *et al*.^32^ Since the lipid phase transition temperature of pure POPC bilayers is *T*_m_ = −2 ^*°*^C,^26^ Subczynski *et al*.’s EPR signals and oxygen transport parameter data represent the data in the gel and fluid phases of POPC membranes, respectively. In Ref.,^32^ the former (EPR signals at *T* = −165 ^*°*^C) suggested that the TEMPO-PC is located in more hydrophilic region, close to the water/lipid interface, than 5-PC or 7-PC, whereas the latter (oxygen transport parameter data at *T* = 25 ^*°*^C) pointed the TEMPO in lipophilic location comparable to 5-PC. These results obtained at two different temperatures comport well with the picture painted by our membrane phase-dependent location of TEMPO moiety.
4. According to a recent experimental study by An *et al*.,^33^ sterol-derived Raman tags introduced in lipid membrane exhibit head-in and head-out types of orientational dimorphism. It was shown that the head-out orientation was preferred when the terminal side chain attached to cholesterol was polar, and that when a terminal side chain was incapable of making hydrogen bonds with water, the head-in configuration was favored at increased temperatures to minimize the perturbation to the lipid-water interface hydrogen bonding (HB) network,^33^ the situation of which is similar to the TEMPO spin label of TEMPO-in configuration submerged in the lipid phase as demonstrated in Fig. 2B. The explanation based on lipid-water hydrogen bond network, given for the orientational dimorphism of sterol-derived Raman tags, however, does not directly apply to the two alternative configurations of TEMPO-PC, given that the average number of H-bonds between TEMPO-PC and water molecules decreases, albeit minor, from TEMPO-out to TEMPO-in configurations (Fig. S3). We surmise that gain in configurational entropy of flexible headgroup and nonpolar interaction between TEMPO moiety and lipid tail (Fig. 2B) is the driving force that gives rise to the TEMPO-in configuration when lipid bilayers are in the fluid phase.

To recapitulate, despite some variations in the PMFs obtained for distinct lipid types and MD force fields (Figs. 3 and 4), our study using MD simulations carried out for single component bilayer membranes leads to the conclusion that the location of the TEMPO moiety of TEMPO-PC in bilayers is membrane phase-dependent, which we hope constructively addresses some of the issues discussed in the field.^15,17,18^ Lastly, given that membrane phase or membrane fluidity depends on a number of factors, such as temperature, types of fatty acids, presence of cholesterol, membrane curvature, medium osmolality, solutes in aqueous phase, and whether the measurement is made on single or multi-bilayer vesicle,^4,34–40^ extra attention should be paid to the physicochemical condition of bilayers when interpreting the data collected from TEMPO-PC.

## METHODS

### Preparation of systems, input files, and analysis

We modeled TEMPO-PC by assembling TEMPO and DPPC using the CHARMM36 force field.^41,42^ Four TEMPO-PCs were placed (two TEMPO-PCs in each leaflet) in DPPC bilayer (64 lipids in each leaflet) and assembled with 150 mM NaCl ions (15 Na^+^ and 15 Cl^*−*^) and SPC water models filling the remaining space of the simulation box. The membrane assembly, necessary input files, forcefield format conversion, and simulation protocols were set up by employing the CHARMM-GUI Membrane Builder.^43,44^ All simulations were performed using GROMACS 2020.4.^45^ The origin of the system was set to the center of *L*_*x*_ × *L*_*y*_ × *L*_*z*_ periodic box (*L*_*x*_ = *L*_*y*_ ≈ 65 Å, *L*_*z*_ ≈ 90 Å). After minimizing the energy of the system using the steepest descent algorithm, the system was equilibrated for ∼ 2 ns by gradually reducing the positional and dihedral restraints on the lipids. In the equilibration step, simulation was carried out in the NVT ensemble with 1 fs integration time step, followed by that in the NPT ensemble with 2 fs time step. The 500 ns production runs in NPT ensemble were generated at *P* = 1 bar and over the temperature range *T* = (260 − 330) K. The semi-isotropic Parrinello-Rahman method and the Nosé-Hoover thermostat were employed to realize the conditions of the constant pressure and temperature. The bond lengths involving hydrogen atoms were constrained using the LINCS algorithm. Area per lipid (*A*), bilayer thickness (*D*_HH_), and density profiles along the bilayer depth were calculated using the last 200 ns of the simulation trajectory. In the NPT ensemble, the *A* was calculated by dividing the area of the simulation box in the *xy* direction with the number of lipids in one of the leaflets (66 lipids; 64 DPPC + 2 TEMPO-PC). In addition, the *D*_HH_ was calculated by taking the average over the *z* values between the phosphorus atoms of the lipids in the upper and lower leaflets.

### Potential of mean force (PMF) of the TEMPO group

The umbrella sampling was carried out to calculate the PMF of TEMPO along the DPPC bilayer at *T* = (290−320) K. To eliminate intermolecular interaction between TEMPO moieties, only a single TEMPOPC was placed in each leaflet. Using the last snapshot of the foregoing MD simulation as the initial configuration, we pulled the oxygen atom of the TEMPO in each leaflet along the *z*-axis using a harmonic potential with stiffness constant *k* = 100 kJ·mol^*−*1^·Å^*−*2^. In total 31 sampling windows with 1 Å-sized bin were placed over 0 Å ≤ |*z*| ≤ 30 Å. After 100 ps equilibration with *k* = 100 kJ·mol^*−*1^·Å^*−*2^ positional restraint, umbrella sampling was conducted for 5 ns at each bin with the umbrella stiffness *k* = 35 kJ·mol^*−*1^·Å^*−*2^ using PLUMED.^46,47^ Weighted histogram analysis method (WHAM) was used to combined the results from all the windows to build the PMF of TEMPO. The position of phosphorus atom within ±10 Å in *xy*-plane from the oxygen radical of TEMPO moiety was set to Δ*z* = 0 of the PMF. Finally, the PMFs were obtained by averaging over all the five replicas.

PMF calculation was also repeated in DOPC and POPC bilayers at *T* = 310 K under CHARMM36^41,42^ and AMBER Lipid17.^48^ For the case of AMBER Lipid17, we adopted the TEMPO model by Stendardo *et al*.^49^

## Acknowledgements

This work was supported by the KIAS Individual Grants CG080501 (S.K.) and CG035003 (C.H.). We thank the Center for Advanced Computation in KIAS for providing computing resources.

## Supplementary Figures

**Figure S1:**
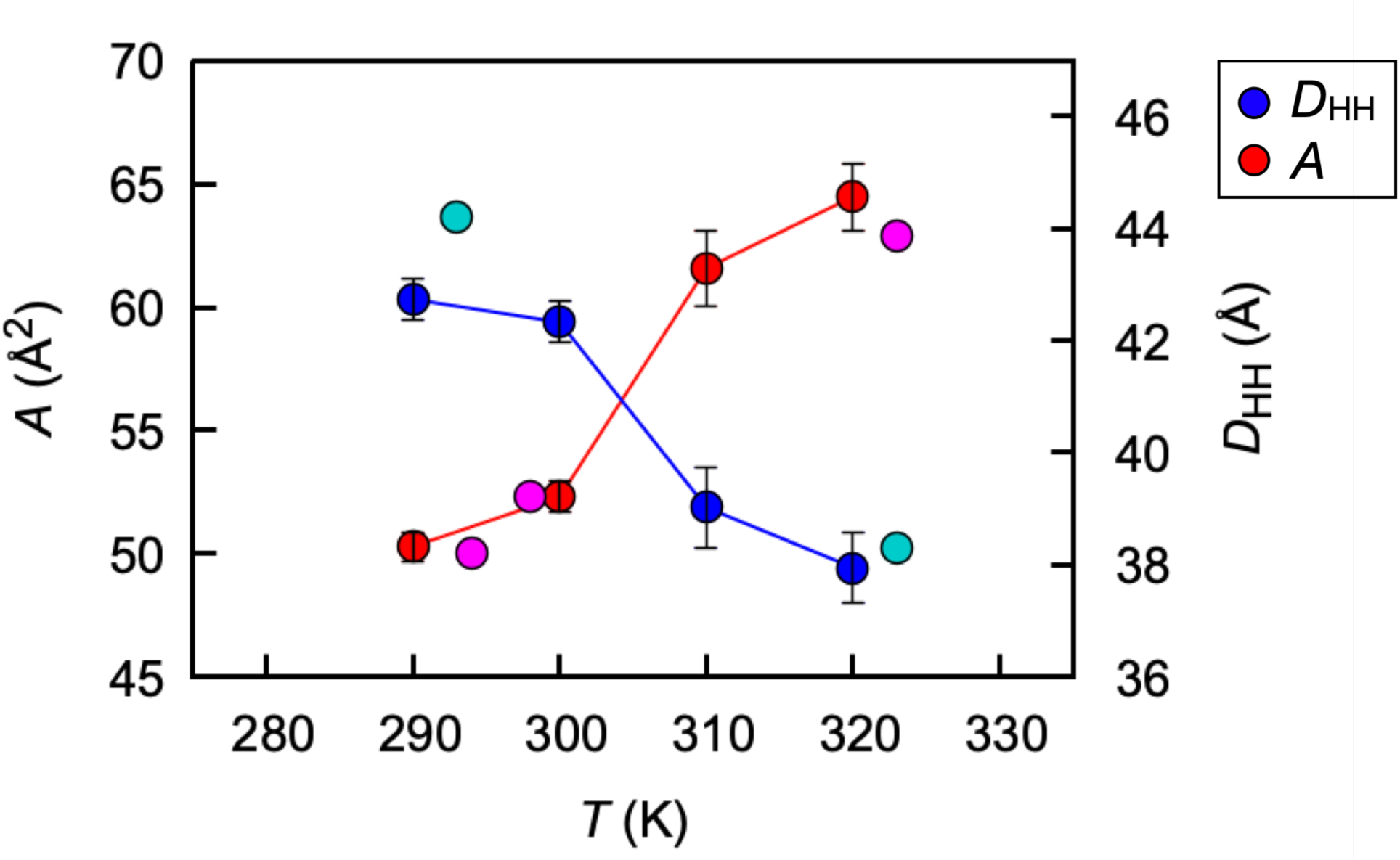
Area per lipid (*A*) and bilayer thickness (*D*_HH_) of DPPC bilayers obtained from MD simulations with AMBER LIPID17 force field at varying temperatures (error bars denote the standard deviation). The filled circles in magenta and cyan represent the experimental measurements for *A* and *D*_HH_, respectively.

**Figure S2:**
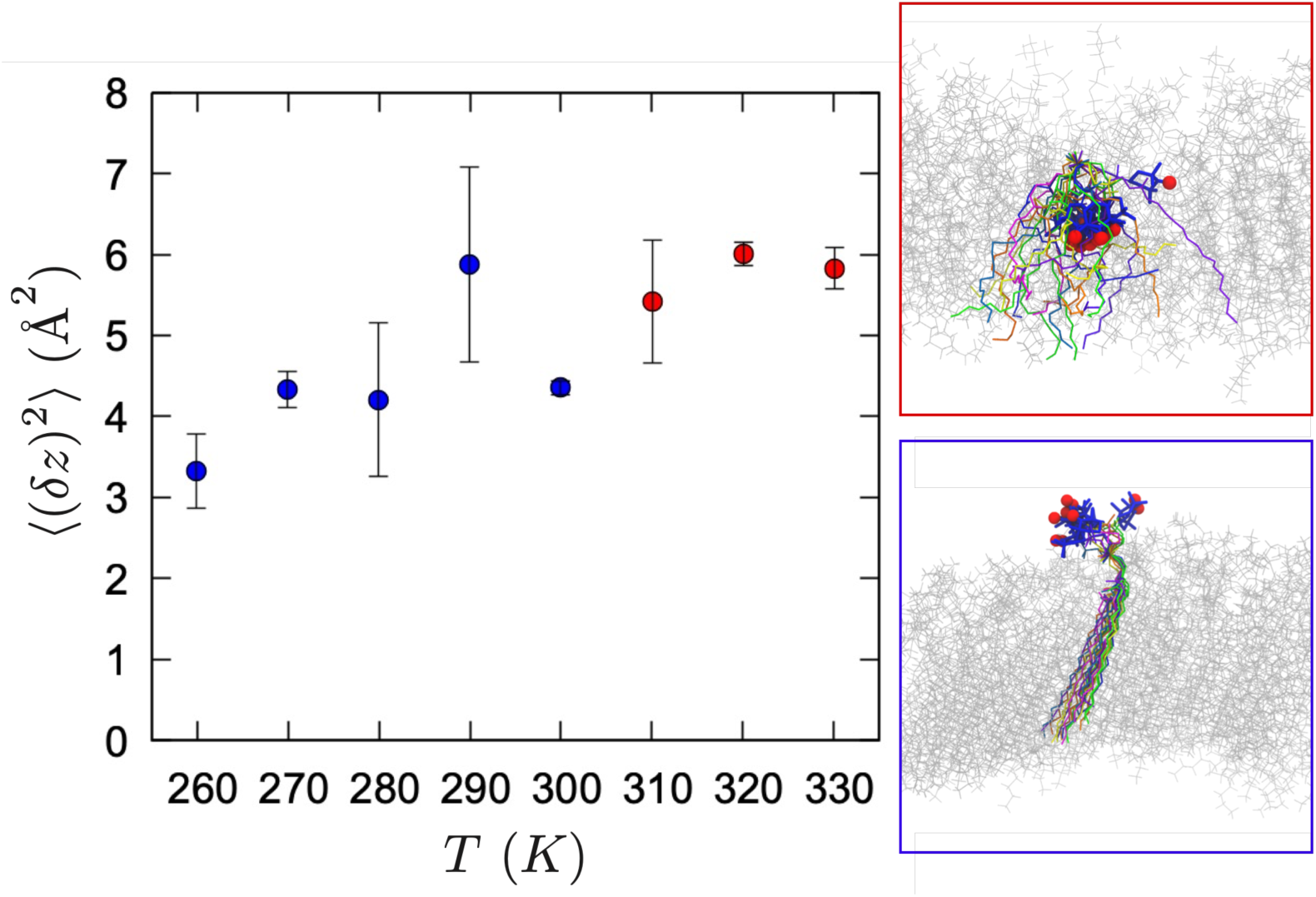
Fluctuations of TEMPO perpendicular to the bilayer plane, ⟨(*δz*)^2^⟩ (variances in the *z*-component of the nitroxide oxygen coordinate), as a function of temperature calculated based on the MD simulation trajectories used to generate the density profiles shown in Fig.1A. The values of ⟨ (*δz*)^2^⟩ in red (*T > T*_*m*_ = 314 K) and in blue (*T < T*_*m*_) in the main panel on the left are calculated by using only TEMPO-in and TEMPO-out configurations, respectively. Two panels on the right depicts the extent of fluctuations in the TEMPO-in at 320 K (red box) and TEMPO-out configurations at 280 K (blue box).

**Figure S3:**
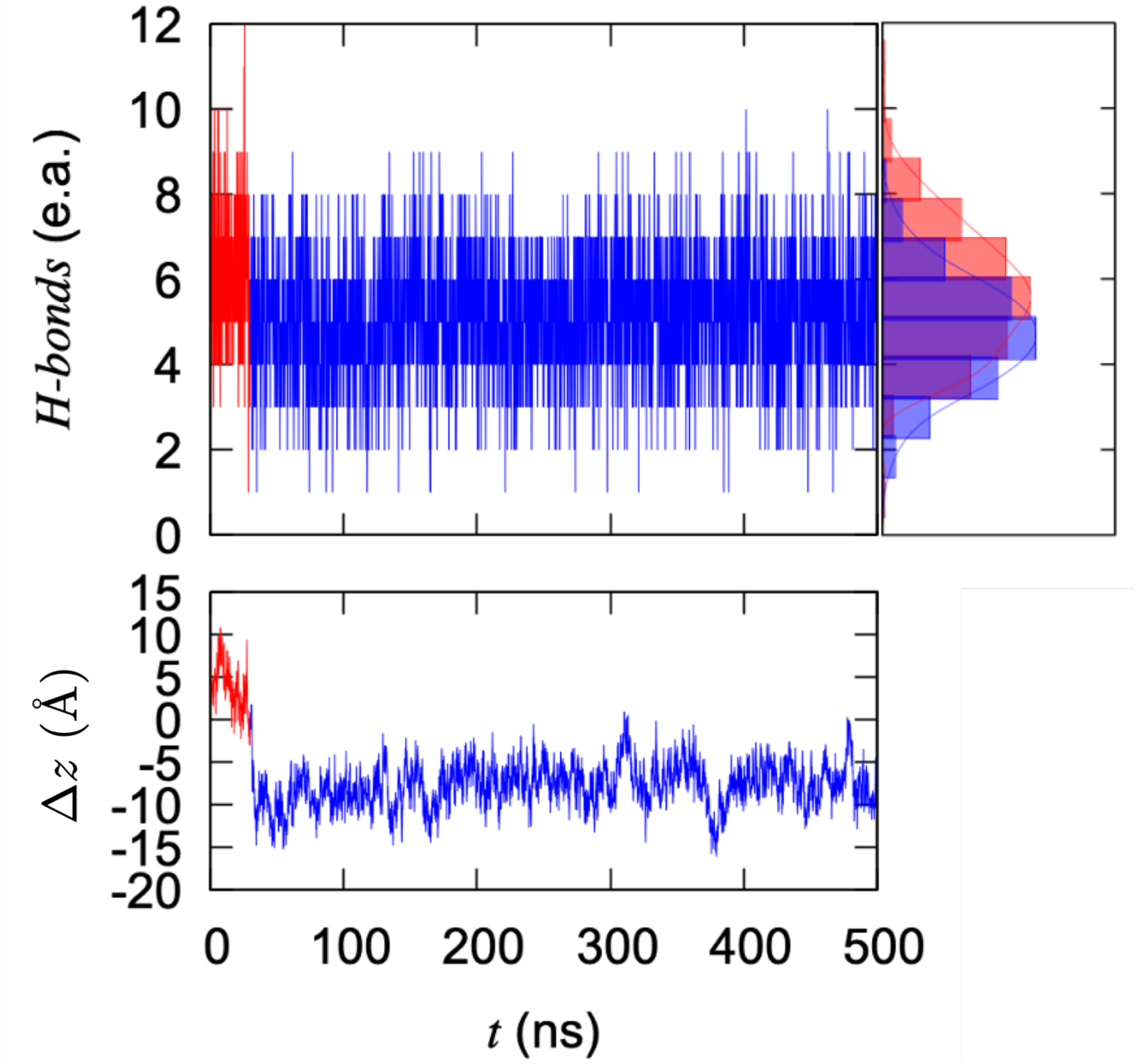
Number of hydrogen bonds between TEMPO-PC and water molecules for the simulation performed at *T* = 320 K. The TEMPO-PC switches its configuration from TEMPO-out (Δ*z >* 0, red) to TEMPO-in (Δ*z <* 0, blue) at *t* ≈ 30 ns. Upon this change, the number of H-bonds decreases from ∼ 6 to ∼ 5.

## Graphical TOC Entry

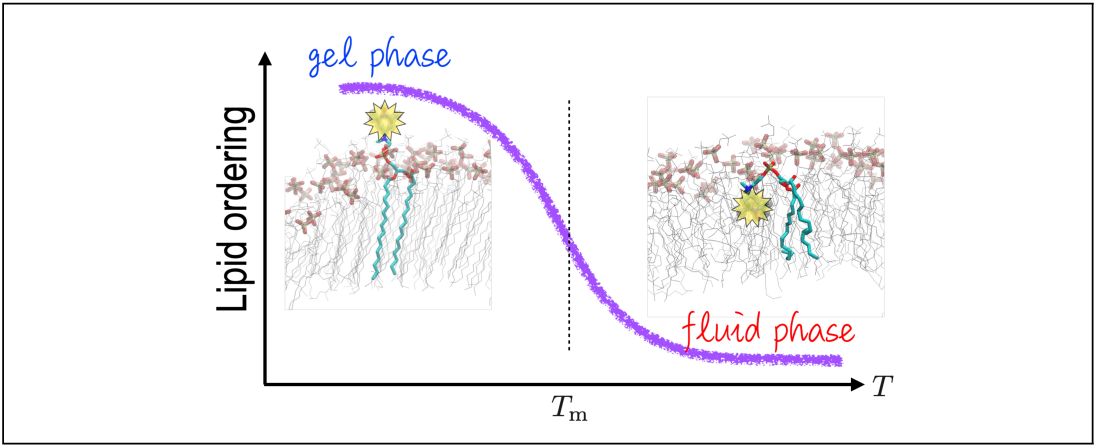

